# A genome-wide ATLAS of liver chromatin accessibility reveals that sex dictates diet-induced nucleosome dynamics

**DOI:** 10.1101/2024.11.13.623052

**Authors:** Zhengyi Zhang, Vivien Su, Carrie B. Wiese, Lijing Cheng, Dan Wang, Ya Cui, Aneesh Kallapur, Jason Kim, Xiaohui Wu, Peter H. Tran, Zhenqi Zhou, David Casero, Wei Li, Andrea L. Hevener, Karen Reue, Tamer Sallam

**Affiliations:** Division of Cardiology, Department of Medicine, University of California, Los Angeles, Los Angeles, California, USA; Department of Physiology, University of California, Los Angeles, Los Angeles, California, USA; Molecular Biology Institute, University of California, Los Angeles, Los Angeles, California, USA; Human Genetics, David Geffen School of Medicine, University of California, Los Angeles, CA, 90095, USA; Division of Computational Biomedicine, Biological Chemistry, University of California, Irvine, Irvine, California, USA; Division of Endocrinology, Diabetes and Hypertension, Department of Medicine, University of California, Los Angeles, Los Angeles, California, USA; F. Widjaja Foundation Inflammatory Bowel & Immunobiology Research Institute, Cedars-Sinai Medical Center, Los Angeles, CA, USA; Department of Medicine and VA Greater Los Angeles Healthcare System GRECC, Los Angeles, California USA 90073; Iris Cantor-UCLA Women’s Health Research Center, Los Angeles, California USA 90095

**Keywords:** chromatin accessibility, sex differences, transcription factor

## Abstract

The three-dimensional organization of the genome plays an important role in cellular function. Alterations between open and closed chromatin states contributes to DNA binding, collaborative transcriptional activities and informs post-transcriptional processing. The liver orchestrates systemic metabolic control and has the ability to mount a rapid adaptive response to environmental challenges. We interrogated the chromatin architecture in liver under different dietary cues. Using ATAC-seq, we mapped over 120,000 nucleosome peaks, revealing a remarkably preserved hepatic chromatin landscape across feeding conditions. Stringent analysis of nucleosome rearrangements in response to diet revealed that sex is the dominant factor segregating changes in chromatin accessibility. A lipid-rich diet led to a more accessible chromatin confirmation at promoter regions in female mice along with enrichment of promoter binding CCAAT-binding domain proteins. Male liver exhibited stronger binding for nutrient sensing nuclear receptors. Integrative analysis with gene expression corroborated a role for chromatin states in informing functional differences in metabolic traits. We distinguished the impact of gonadal sex and chromosomal sex as determinants of chromatin modulation by diet using the Four Core Genotypes mouse model. Our data provide mechanistic evidence underlying the regulation for the critical sex-dimorphic GWAS gene, *Pnpla3*. In summary, we provide a comprehensive epigenetic resource in murine liver that uncovers the complexity of chromatin dynamics in response to diet and sex.

**Highlights:**

ATAC-Seq, RNA-Seq, and FCG model-integrated analysis unravel sex differences in chromatin accessibility and transcriptome responses to dietary challenges.
Lipid-rich diet led to sex-biased chromatin confirmation at promoter regions.
Gonadal sex emerged as the most prevalent determinant of the sex bias hepatic chromatin modulation by lipid-rich diets.
The critical sex-dimorphic GWAS gene *Pnpla3* is suppressed by testosterone, which underlies hepatic differences in expression between the sexes.

## Introduction

An important way that cells control transcript biogenesis and crosstalk with other steps in gene regulation is through epigenetic checkpoints (Kan et al., 2022). DNA is typically stored in highly condensed structures known as nucleosomes which then congregate to form chromatin (Maeshima et al., 2021). Chromatin can convert from an active to inactive state through a variety of mechanisms (Klemm et al., 2019). Chemical modifications on DNA or histone proteins, for example, can alter the three-dimensional chromatin structure and nucleosome positioning (Becker and Workman, 2013; Garcia-Ramirez et al., 1995). Since rearrangements in nucleosomes often precede transcription factor binding and activation (Barozzi et al., 2014; Suter, 2020), information about nucleosome position can be used to infer active regulatory regions. Genome-wide interrogation of accessible chromatin regions grew in popularity with the development of ATAC-seq (Buenrostro et al., 2015; Grandi et al., 2022). This robust technique leverages a Tn5 enzyme that permits the determination of a genome-wide epigenetic map. Despite an increase in ATAC-seq datasets that describe spatial organization of chromatin states in tissues under basal conditions or during development (Deng et al., 2022; Liu et al., 2019; Trevino et al., 2020), our understanding of how chromatin rearrangements inform metabolic health and disease states remains limited. In particular, few studies have carefully examined sex-differences in nucleosome positioning in response to dietary cues.

The liver is an important metabolic center that transduces signals from diet to influence gene expression and metabolism (Bideyan et al., 2021; Yang et al., 2006). For example, high fat diet feeding imparts unique perturbations in liver metabolism to accommodate the intake of excess calories and high fat content. Similarly, a diet rich in cholesterol leads to changes in liver metabolism and RNA biogenesis that is distinct from high fat diet feeding (Hui et al., 2015; Salisbury et al., 2021; Soltis et al., 2017; Xiao et al., 2023). Both high fat and cholesterol-rich diets can increase the risk of Metabolic Associated Fatty Liver Disease (MAFLD) (Gallage et al., 2022; Im et al., 2021; Radhakrishnan et al., 2021).

It is well established that many metabolic traits, including MAFLD, are influenced by biological sex (Lonardo et al., 2019; Meda et al., 2020; Pan and Fallon, 2014). For example, there are known sex-biased differences in hepatic lipid composition at baseline and following feeding lipid-rich diets (Meda et al., 2020; Salisbury et al., 2021). Mediators that are proposed to confer sex differences in hepatic gene expression and metabolism include the estrogen receptors, growth hormone, X-chromosome factors, and the BCL6-STAT5 axis (Blencowe et al., 2022; Salisbury et al., 2021). Relatively little is known about the interaction between sex and diet at the level of chromatin organization and transcriptional dynamics. Here, we present an unbiased analysis of transcriptional regulation by diet and sex determinants, which has uncovered previously unappreciated relationships.

One level of control by which the liver responds to different challenges is by regulating the chromatin landscape. Changes in chromatin can lay the foundation for transcription or interface with pathways involved in transcript splicing, modification, and export (Li et al., 2007; Petrova et al., 2022; Simon et al., 2014; Xiao et al., 2019; Yoh et al., 2007). To what extent does the chromatin environment in the liver change with different cues, and how does it contribute to gene expression and function? In this study, we map the chromatin profile in mouse liver and show that sex primes the chromatin landscape in liver and biases responses to lipid-rich diet. Using an example a of disease-relevant gene, *Pnpla3*, we reveal the molecular underpinning of sex-differences in MAFLD that consolidates well with known clinical observations.

## Results

### Chromatin accessibility profiling in liver is dictated by sex at promoter regions

To better understand how chromatin rearrangements are informed by diet composition we performed ATAC-seq in liver of male and female mice fed with normal chow, western diet (WD, rich in cholesterol) or high fat diet (HFD, low cholesterol but rich in fat content) for 2 weeks (Figure 1A). We obtained comparable read counts across all groups after sequencing, and each sample exhibited a similar alignment rate to the mouse genome mm9 (Figure S1A and S1B).

**Figure 1.**
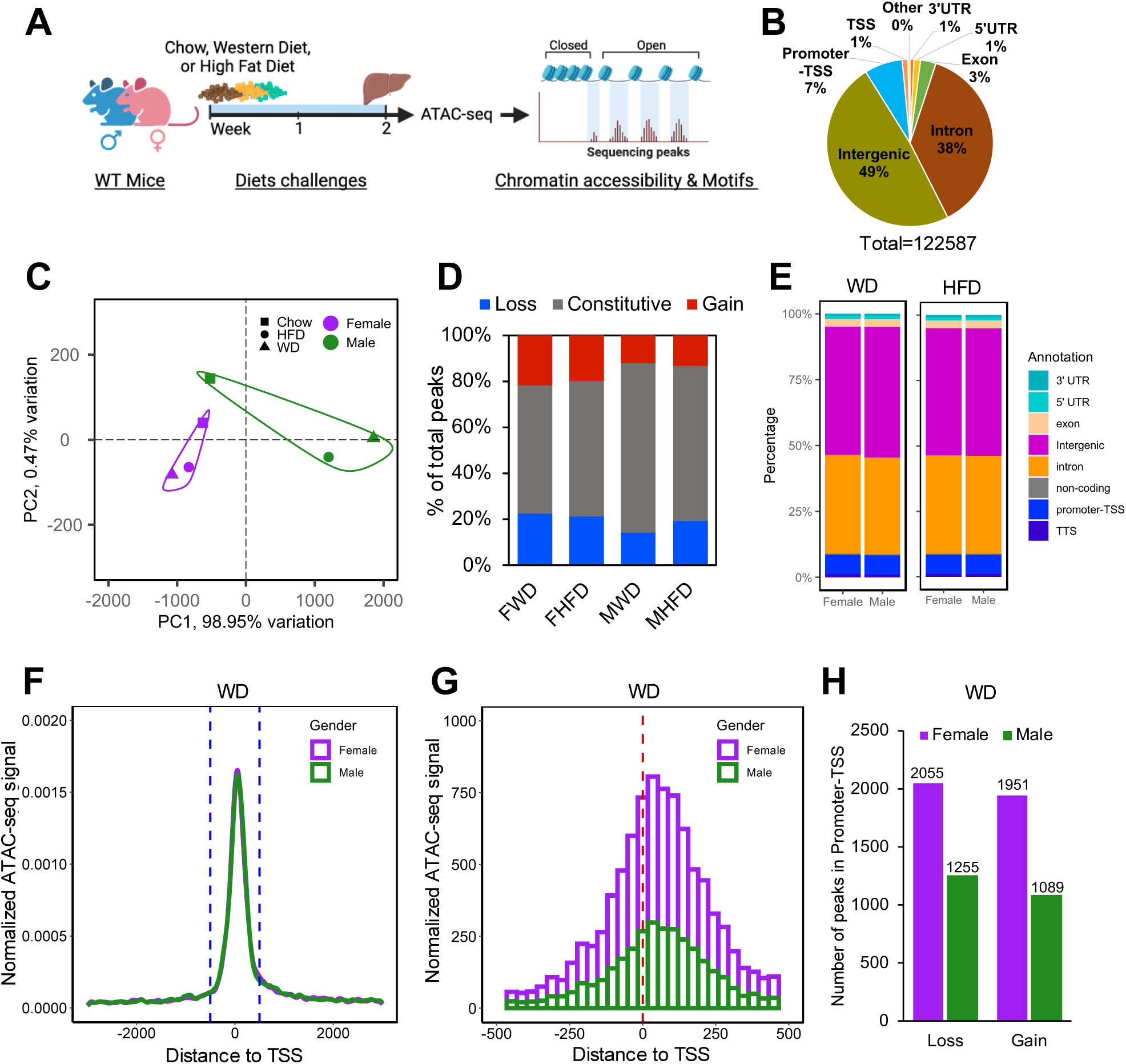
Sex segregates liver chromatin accessibility in response to diet. (A) Design of liver ATAC-seq experiments (N= 3 mice per group). (B) Distribution of genomic features for all peaks. (C) PCA plots of chromatin accessibility of liver samples using mean peak score from each group (n=122,587). (D) Comparison of the proportion of genes. FWD: female WD peaks compared to female chow peaks; FHFD: female HFD peaks compared to female chow peaks; MWD: male WD peaks compared to male chow peaks; MHFD: male HFD peaks compared to male chow peaks. Peaks with fold change (FC)> 2 were considered ‘gain of accessibility’ and FC< 0.5 were considered ‘loss of accessibility’. (E) Distribution of genomic features amount differential peaks (FC> 2 or FC< 0.5) in female or male mice liver. (F) and (G) Normalized ATAC-seq signal intensity for differential peaks of WD compared to chow ± 3000 nt (F) or ± 500 nt (G) from TSS. (H) Number of differential female and male peaks in the Promoter-TSS region according to Homer annotation. Chow: chow diet; WD: western diet; HFD: high fat diet.

Across conditions, we mapped a total of 122,587 unique peaks (Figure 1B) (Table S1), the majority of which are located in Intergenic (49 %) or Intronic DNA regions (38 %), with 7 % of peaks located in “promoter-TSS” regions (Figure 1B). Principal component analysis–a dimensionality technique that explores contributors to variance–of liver ATAC-seq showed that chow-fed female and male had largely similar profiles for principal component PC1 and PC2 (Figure 1C and Figure S1C). PC1 captured >98% of variance observed across conditions (Figure 1C). Compared to females, male mice exhibited greater variability, particularly when subjected to WD or HFD challenges (Figure S1C). Interestingly, WD or HFD feeding induced completely distinct responses on chromatin landscape with respect to sex. Feeding of HFD or WD further segregated changes in chromatin landscape with WD leading to more dramatic differences across PC1. This result revealed that hepatic chromatin accessibility differences in response to dietary lipid excess are orchestrated in a highly sex-biased manner.

While most peaks did not change access in response to diet (constitutive peaks), females exhibited both more loss and gain of accessible chromatin sites compared to males (Figure 1D). The percentage of differential chromatin changes (DCC) resulting from WD and HFD did not reveal notable sex-specific differences in genomic localization (Figure 1E). Peak distribution analysis revealed that sex-induced chromatin differences cluster around the TSS region, with comparable distribution for both sexes in other regions (Figure 1F, G, Figure S1D and S1E). Compared to chow baseline, WD-fed or HFD-fed female mice showed more DCCs, for both gain and loss of access sites, at promoters compared to WD-fed or HFD-fed male mice (Figure 1H, Figure S1F). Collectively, our findings suggest male and female mice exhibit distinct changes in chromatin rearrangements, particularly at promotor regions, in response to a hyperlipidemic diet.

### Motif analysis reveals a sex-biased TF preference in response to diet challenges

We next analyzed motif preferences using the DCCs from male and female mice to decipher the transcription factors involved in the sex-specific chromatin regulation. Since sex bias was noted in chromatin accessibly at promoter regions, we ranked all promoter-specific differential peaks based on changes in accessibility (HFD vs chow diet, or WD vs chow diet) in both sexes and binned them into equal-sized bins (six bins each), followed by motif enrichment analysis (see Methods for details). This strategy allows us to compare the enrichment of each motif across all bins using a significance enrichment score (Figure S2A). While most motifs exhibited similar enrichment between males and females, some motifs showed a sex-biased enrichment pattern for certain bins or groups of genes, including ETS family transcription factors (TFs) (Elk4, ETS1, EVT4, GABPA) as well as the promoter-binding transcription factor NFY (Figure S2A). While enriched in both male and female mice, NFY showed differential enrichment between males and females in the HFD and WD group (Figure S2B). Among the top 10 enriched motifs from each group, 9 were found to be commonly enriched (Figure S2C). While stronger enrichment was observed for certain transcription factors in males (KLF3, KLF5, Elk1, Elk4, and Sp5), NFY emerged as a strongly enriched transcription factor in females (Figure S2D). Collectively, these results suggest that binding patterns of most TFs at promotor regions respond similarly to diet in both sexes, but there is sex bias in some groups of TFs, including NFY, that are known to be directly involved in regulating chromatin access and collaborative TF binding.

To further disentangle the factors that led to sex-specific chromatin responses to hyperlipidemia at a genome-wide scale, we separated signatures of male-biased and female-biased DCCs (Figure 2A): C1 and C8 quadrants (C1&C8) represent male-specific enhanced chromatin opening in response to WD (group I); C5&C6 represent the female-specific enhanced chromatin opening in response to WD (group II); C4&C5 represent the male-specific reduced chromatin opening in response to WD (group III), and C1&C2 represent the female-specific reduced chromatin opening in response to WD (group IV). Motif analysis of the top or bottom 100 peaks for each sex (Figure 2B) revealed distinct positional bias for each group (Figure 2C). For example, Cux2 was specifically enriched in “female loss” of accessibility cluster (group IV), consistent with previous reports showing that Cux2 tends to be more enriched at the genomic region exhibiting male-higher accessible chromatin than in female mouse liver (Ling et al., 2010). Female-biased clusters also showed enrichment of estrogen receptor motifs, as expected. The “male gain” (group I) and “female loss” (group IV) peaks preferentially exhibited enrichment of motifs of well-known nutrient-sensing nuclear receptor family members such as LXRE, RXRE and PPARE, and the “female gain” (group II) and “male loss” (group III) peaks specifically enriched the motif of Zinc Finger family (Zf), ETS family and NFY (CCAAT) (Figure 2C). Notably, the top enriched motif for “male gain” of chromatin access is LXRE while the top enriched motif for “female gain” of chromatin access is NFY (Figure 2D).

**Figure 2.**
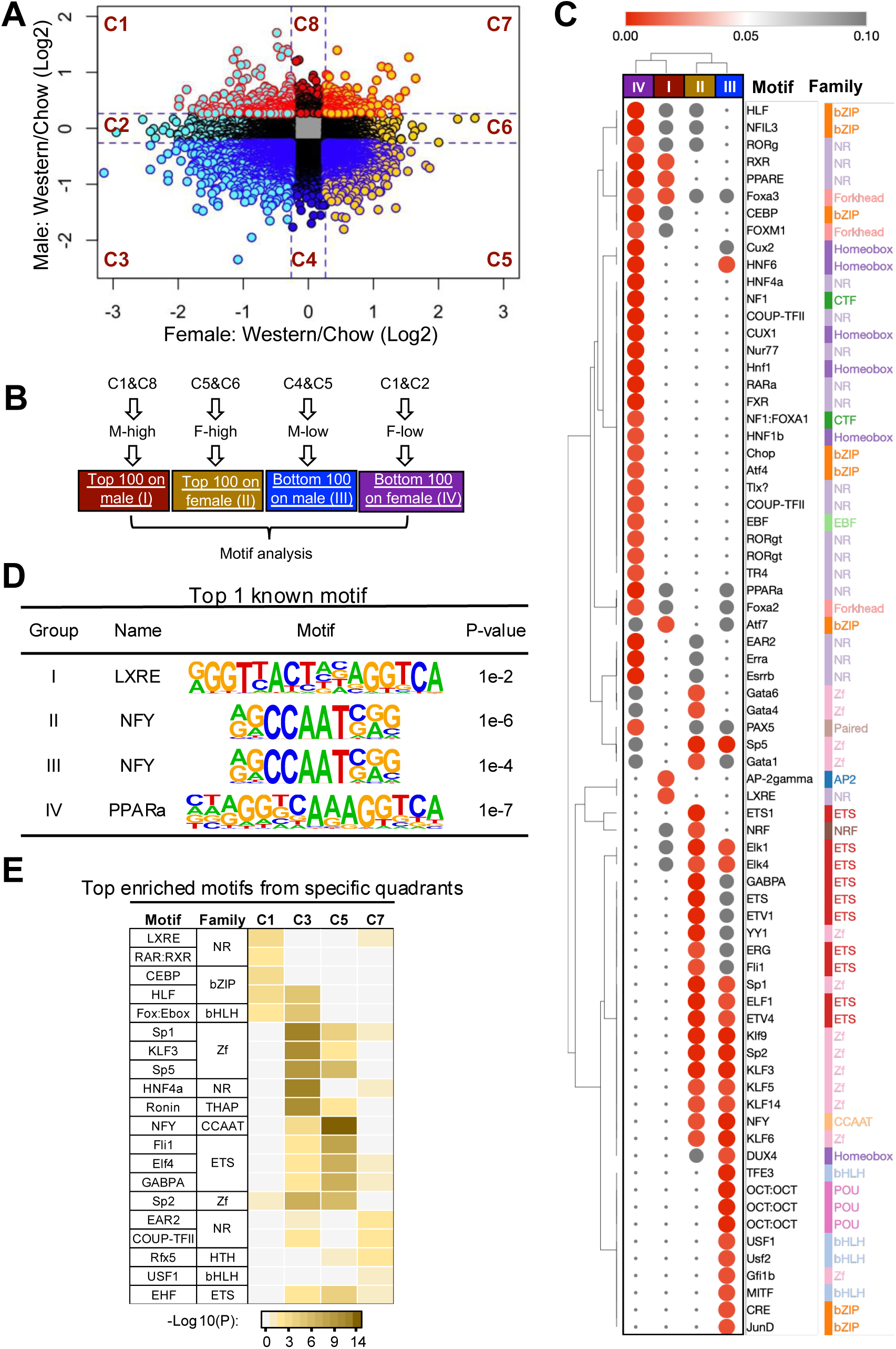
Diet induces sex-divergent motif enrichment in the liver. (A) Correlation analysis of male and female ATAC-seq peaks under WD. Blue lines indicate 1.2-fold change. (B) Motif analysis group strategy. (C) Enriched motifs and motif families in different group in (B) as analyzed by Homer. Color in heatmap represents the P-value. (D) Top 1 motif from each group from (C). (E) Top 5 enriched motifs from specific quadrants and their enriched P-value from other quadrants.

To focus on the motifs that show the greatest sex bias, we reanalyzed the motifs using quadrants C1, C3, C5 and C7 in Figure 2A, representing concordant sex responses (“male & female gain” and “male & female loss”) or divergent sex responses (“male gain & female loss” and “female gain & male loss”). Once again, the results demonstrate that LXRE was more strongly enriched in “male gain & female loss” group (C1), while NFY was preferentially enriched in the “female gain & male loss” group (C5) (Figure 2E). Taken together, these results hint that in response to diet feeding, males may be more sensitive to chromatin rearrangements involving nutrient-sensing transcription factors whereas females are more sensitive to promotor-binding factors such as NFY.

### Nucleosome dynamics of sex-specific transcription factors are associated with distinct transcriptomic responses

To gain insight into the consequences of the hepatic chromatin accessibility/TF bias between males and females in driving differential gene expression, we determined the hepatic transcriptome differences between male and female mice responding to 2 or 4 weeks of WD or HFD feeding compared to chow (Figure 3A). Consistent with ATAC-seq results, RNA-seq revealed that WD and HFD induced or suppressed gene expression in similar patterns (Figure S3A). While the majority of genes were unchanged with a lipid-rich challenge, there were approximately 2,000 differentially expressed genes (DEGs) in response to WD and HFD (Figure S3B and S3C) (Table S2). The vast majority of these DEGs (∼80%) exhibited sex-biased expression based on fold changes and P-values (described in Materials and Methods) (Figure 3B and 3C). This observation is in line with other evidence showing that a significant proportion of hepatic genes show sex-biased regulation (Yang et al., 2006). To decipher sex-specific DEGs, we compared the female and male DEGs from 2 and 4 weeks lipid-rich dietary challenge (Figure S3D) and functionally annotated these genes by Metascape analysis (Zhou et al., 2019). Shared DEGs between males and females showed “Metabolism of lipid” as the top enriched pathway (Figure S3E). Female- and male-specific DEGs in response to diet exhibited differences in pathway enrichment. Male DEGs showed enrichment of metabolic pathway changes whereas female DEGs showed stronger changes in cell migration and MCM (minichromosome maintenance) remodeling (Figure S3F-I).

**Figure 3.**
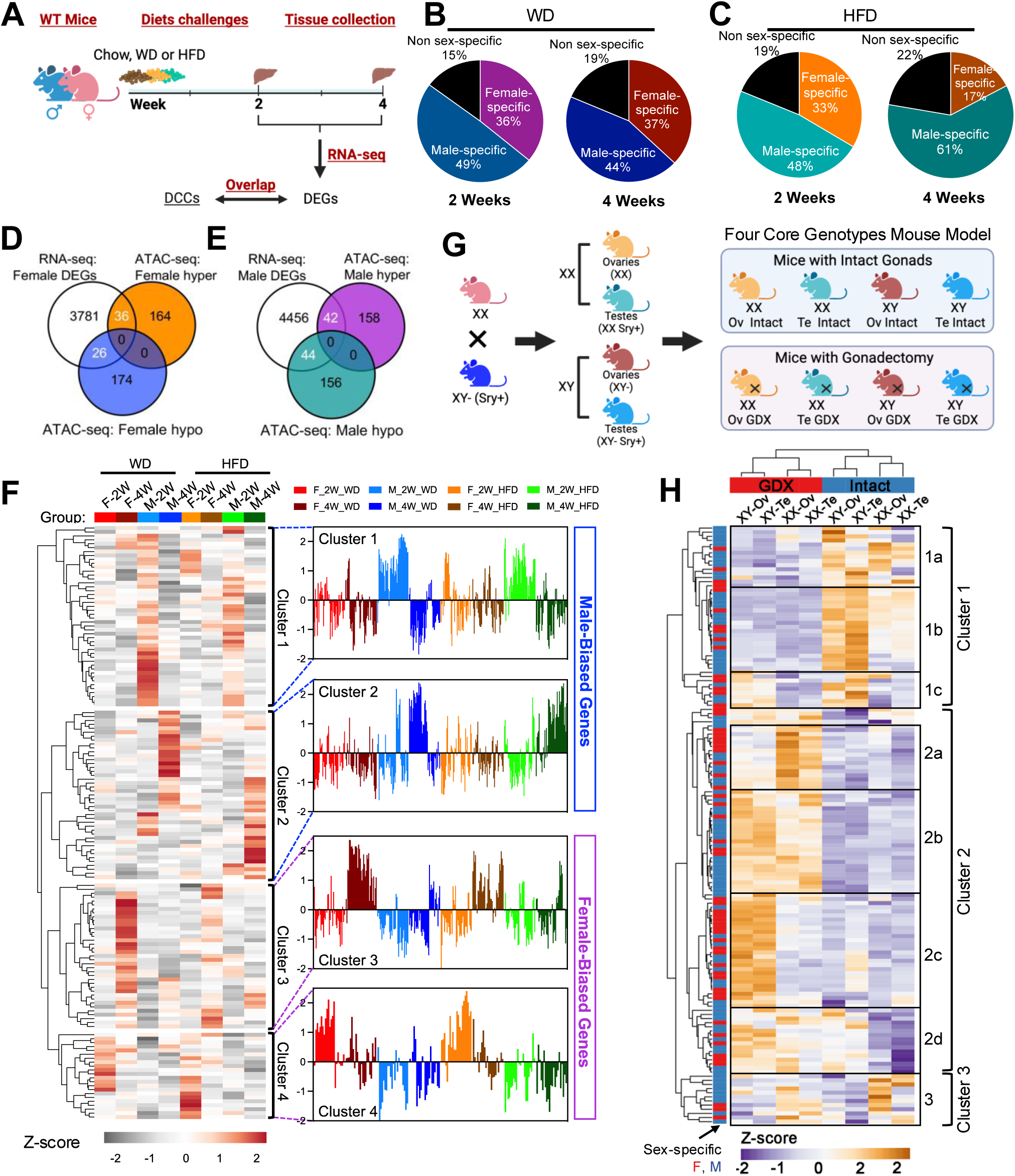
Integrative analysis identifies sex-dimorphic chromatin-gene expression connections in liver tissue. (A) Schematic showing experiment design of RNA-seq in mice liver with diet feeding. (B) and (C) Differentially expressed genes (DEGs) in male or female with 2 weeks or 4 weeks of WD (B) or HFD feeding (C). Cutoff: |FC| > 1.5 & P< 0.05. (D) DEGs of female in 2 weeks and 4 weeks overlapped with “female hypo” ATAC-seq peaks (Bottom 200 peaks based on female in C1&C2 clusters of Figure 2A) or “female hyper” ATAC-seq peaks (Top 200 peaks based on female in C5&C6 clusters of Figure 2A). (E) DEGs of male in 2 weeks and 4 weeks overlapped with “male hypo” ATAC-seq peaks (Bottom 200 peaks based on male in C4&C5 clusters of Figure 2A) or “male hyper” ATAC-seq peaks (Top 200 peaks based on male in C1&C8 clusters of Figure 2A). (F) Heatmap was generated based on fold changes calculated by RNA-seq RPKM of overlapped genes from Figure 3D and Figure 3E. F-2W: female fold changes calculated by WD compared to Chow or HFD compared to Chow at 2-week time point; F-4W: female fold changes calculated by WD compared to Chow or HFD compared to Chow at 4-week time point; M-2W: male fold changes calculated by WD compared to Chow or HFD compared to Chow at 2-week time point; M-4W: male fold changes calculated by WD compared to Chow or HFD compared to Chow at 4-week time point; (G) Schematic of the Four Core Genotypes (FCG) mouse model. (H) The expression of diet-induced, sex-biased genes, as shown in Figure 3F, was detected using RNA-seq in the FCG mouse model in (G). Clusters were generated based on CPM. Ov: ovaries, Te: ovaries.

To explore the relationship of genes with sex-specific differential expression to chromatin accessibility changes, we overlapped DEGs from male or female RNA-seq with female or male ATAC-seq changes (Figure 3D and 3E). Clustering of gene expression by integrating ATAC-seq showed that WD and HFD feeding have similar overall effects on gene expression (Figure 3F). However, both WD and HF feeding showed divergent gene expression patterns with respect to feeding time (Figure 3F). Furthermore, based on gene expression patterns, we predicted the top enriched motifs of male-biased and female-biased genes (Figure S3J). Our analysis identified enrichment of androgen receptor (AR) motif among male-biased genes with low enrichment in females. Reciprocally, estrogen receptor (ESR1) was strongly enriched among female-biased genes but not males. These results align well with known hormonal regulation of hepatic gene expression in the two sexes. Unlike our ATAC-seq analysis, which showed higher LXR enrichment in males (Figure 2E), LXR was enriched in both male and female clusters based on transcriptomics (Figure S3J). Thus, undertaking an RNA-centric analysis strategy reinforced most but not all of the sex-biased results of chromatin dynamics. These results hint that additional layers in gene regulation beyond transcription and epigenetic control (e.g., transcript stability, export, and processing) may contribute to steady-state transcriptome levels.

### Sex-biased diet responsive genes are influenced by gonadal and chromosomal sex

There are two primary determinants of biological sex—sex chromosomes (XX or XY) and gonads (ovaries or testes and hormones secreted by these tissues) (Lopez-Lee et al., 2024; Mauvais-Jarvis, 2024; Sandovici et al., 2022). To further investigate the role of chromosomal and gonadal sex determinants in sex-biased gene expression in response to a lipid-rich diet, we employed the Four Core Genotypes (FCG) mouse model, which includes four sex genotypes (XX with ovaries, XX with testes, XY with ovaries, XY with testes) (Figure 3G) (Arnold, 2020; Blencowe et al., 2022; Mauvais-Jarvis et al., 2017; Reue and Wiese, 2022; Zhang et al., 2024). We assessed the determinants of gene expression for the 148 sex-biased genes shown in Figure 3F in FCG mice having intact gonads or after surgical removal of gonads (GDX) from adult mice followed by HFD feeding. About 80% genes that responded to lipid-enriched diets in a sex-biased manner in standard male and female mice were also regulated by a sex determinant (gonadal sex, chromosomal sex, or both) in the FCG model (Figure S3K and S3L) (Table S3).

Gonadal status emerged as the most prevalent determinant of the sex bias, although many genes were influenced by both gonadal and chromosomal sex (Figure 3H, Figure S3L, Table S3). Cluster 1 included genes whose expression was diminished when acute gonadal hormones were depleted by removal of gonads, while Cluster 2 consisted of genes inhibited by gonadal secretions, as their expression was enhanced following GDX (Figure 3H). Within these groups, many genes were influenced by chromosomal sex, with clear patterns of greater expression in XX compared to XY liver (for example, cluster 2a, XX>XY in GDX mice), or greater expression in XY compared to XX liver (for example, cluster 1b in intact mice and cluster 2c in GDX mice). These findings reveal the complex regulation of sex-biased diet response genes by both genetic and hormonal sex determinants. In many cases, the regulation by sex chromosome genotype is more apparent after gonadal hormones are removed, suggesting that sex chromosome complement may be an important determinant of diet response under conditions of reduced gonadal hormone levels, such as following menopause and andropause in humans. In summary, integrating hepatic ATAC-seq and RNA-seq under different conditions shows overlap in sex-biased signatures and suggests, at least in part, changes in chromatin dynamics under different conditions are functionally relevant and inform gene expression changes.

### Chromatin architecture informs mechanisms of sex differences at disease relevant genes

To more carefully interrogate how diet composition related to sex-differences in gene expression we stringently defined female-biased and male-biased genes as outlined in our methods. We ultimately identified 25 female-biased DEGs and 32 male-biased DEGs (Figure 4A). Many of these genes showed sex-biased expression in human liver (Figure S4A-S4C) (Table S4). Notably, Patatin-like phospholipase domain-containing protein 3 (*Pnpla3*) emerged as one of the top female-biased in both mouse and human liver (Figure 4A, Figure S4B and S4D). Hepatic *PNPLA3* is of substantial interest because variants at *PNPLA3* have been associated with risk of MAFLD (Anstee et al., 2020; Ma et al., 2017; Romeo et al., 2008; Speliotes et al., 2010). We obtained consistent results by analyzing public RNA-sequencing datasets (Lau-Corona et al., 2020), which suggested that *Pnpla3* is one of the top-ranked female-biased genes (Figure 4B). We confirmed the female-biased expression of *Pnpla3* by qPCR on liver samples from mice fed different diets (Figure 4C). Notably, 16 of the female-biased genes we identified overlapped with those from previously published RNA-sequencing datasets (Lau-Corona et al., 2020), with *Pnpla3* showing the greatest difference between males and females (Figure S4E and S4F).

**Figure 4.**
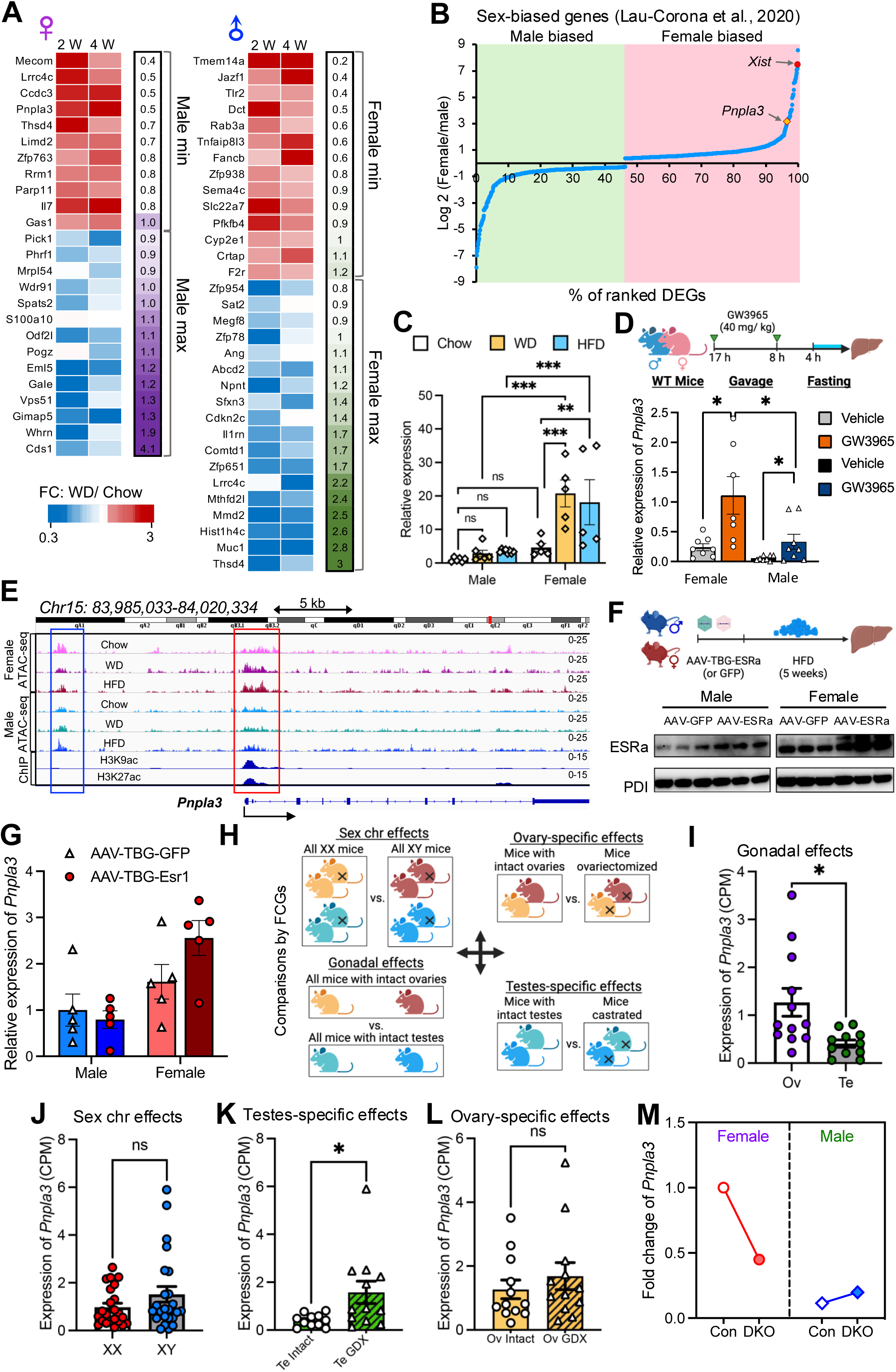
*Pnpla3* is regulated by male gonads and epigenetic factors. (A) Fold changes of potential sex-biased genes are displayed in the heatmap. The left panel shows female-biased DEGs as described in methods, while the right panel presents male-biased DEGs calculated using the same method as the female-biased DEGs. (B) Hepatic sex-specific DEGs were identified by RNA sequencing from Lau-Corona et al., 2020 (detailed in Methods). (C) Gene expression of *Pnpla3* detected by qPCR from livers of mice fed with chow, WD or HFD for 2 weeks. n = 6 (Male Chow and male WD); n = 8 (male HFD); n=5 (Female Chow, WD and HFD). Data are represented as mean ± SEM. P-values were calculated by two-way ANOVA. **: P< 0.01; ***: P< 0.001; ns: not significant. (D) LXR agonist treatment and gene expression of *Pnpla3* detected by qPCR. n = 8 (Female Vehicle, male Vehicle and male GW3965); n = 7 (Female GW3965). *: P< 0.05 calculated by unpaired t test. (E) Peaks of ATAC-seq from this study and ChIP-seq of histone mark from ENCODE project with GEO number: GSM1000153 (H3K9ac); GSM1000140 (H3K27ac). ATAC-seq peaks were overlayed from all 3 samples of each group. (F) Schematic of ESRa overexpression by AAV and immunoblotting of ESRa in murine livers. (G) Gene expression of *Pnpla3* detected by qPCR from livers of mice from Figure 4F. N= 5; not significant by either t-test or ANOVA. (H) Schematic comparing *Pnpla3* expression using the FCG mouse model. (I) Hepatic *Pnpla3* expression in mice with intact ovaries (Ov) or with intact testes (Te). N=12 (Ov), N=11 (Te). *: P< 0.05 by unpaired t-test. (J) Hepatic *Pnpla3* level in mice with chromosome XX and XY. N= 23 (XX). N= 24 (XY). ns: P> 0.05 by unpaired t-test. (K) Hepatic *Pnpla3* level in mice with intact testes (Te Intact) or with testes GDX (Te GDX). N= 11 (Te Intact). N= 12 (Te GDX). *: P< 0.05 by unpaired t-test. (L)Hepatic *Pnpla3* level in mice with intact ovary (Ov Intact) or ovary GDX (Ov GDX). N= 12. ns: P> 0.05 by unpaired t-test. (M) Fold change calculated using FPKM of *Pnpla3* from Lau-Corona et al., 2020. Data shown as FPKM relative to Female control. DKO: Ezh1 and Ezh2 double-KO.

*Pnpla3* is known to be impacted by LXR, and our analysis showed that LXR chromatin dynamics differs between the sexes, although these LXR changes minimally inform global transcriptome signatures. To carefully examine a role for LXR in underlying sex-differences in hepatic *Pnpla3*, we employed a variety of approaches. We assessed changes in canonical LXR target genes during physiologic perturbations. RNA-seq under different dietary conditions showed no major sex differences in the expression of LXR target genes except for *Scd1* under certain conditions (Figure S4G). In addition, we orally administered the LXR agonist GW3965 to male and female mice and examined gene expression by qRCR. LXR targets were induced with GW3965, but we did not observe substantial differences in LXR gene regulation comparing male and female liver (Figure S4H). Collectively, these results suggest that there may be subtle differences at select LXR targets, but response to LXR agonist showed no major ‘global’ sex effect (Figure S4G). However, even with maximal LXR activation, *Pnpla3* was confirmed to be a female-biased gene by qPCR (Figure 4D). Collectively, our data suggest that LXR regulates *Pnpla3* expression, but is not a critical factor in sex-differences in its expression.

Prior work reported that PNPLA3 underlies sex-differences in MAFLD (Kosters et al., 2013). A recent study reported higher hepatic expression of *PNPLA3* in female liver may be driven by estrogen receptor-α (ER-α) (Cherubini et al., 2023). We examined the chromatin architecture at the *Pnpla3* locus in male and female liver. We identified a cluster of accessible regions at the *Pnpla3* promoter region extending into the gene body (red, Figure 4E) and another upstream region (blue). The upstream region (blue) was near estrogen receptor binding sites and showed minimal regulation between the sexes. This led us to reexamine a role for estrogen receptor in dictating the sex bias in *Pnpla3* expression. We treated wild-type male and female mice with an adeno-associated virus (AAV) under control of a TBG promoter, expressing either ER-α (AAV-TBG-ESRa) or GFP (AAV-TBG-GFP). We confirmed robust overexpression by immunoblotting as well as the enhanced expression of *Polg1*, an ER-α regulated gene we previously demonstrated (Figure 4F, Figure S5A). *Pnpla3* expression was measured by qPCR, and no significant difference was observed between AAV-GFP and AAV-ESRa in either males or females (Figure 4G), hinting that estrogen may not be the dominant factor contributing to sex-dependent differences in the regulation of *Pnpla3*. We evaluated *Pnpla3* expression using the FCG mouse model, comparing the effects of sex chromosomes, gonadal status, and ovary/testes-specific influences (Figure 4H). Expression levels were higher in the genotypes with ovaries compared to those with testes (Figure 4I), with no significant difference between the XX chromosome and XY chromosome groups (Figure 4J). We further investigated which gonadal factor regulated *Pnpla3* expression. Removal of testes led to increased *Pnpla3* expression (Figure 4K), whereas removal of ovaries had no effect. (Figure 4L). Overall, these results suggested that *Pnpla3* was suppressed by testosterone rather than regulated by ovarian hormones.

The accessible region at promotors at *Pnpla3* showed strong regulation between male and females fed different diets (Figure 4E, red). Notably, accessibility at the *Pnpla3* promoter was dramatically reduced in male mice (Figure 4E). PROMO (Farre et al., 2003; Messeguer et al., 2002), a method that constructs positional weight matrices from known transcription factor binding sites for a given gene, supports the idea there are potential hotspots around the promoter region of *Pnpla3* that may underlie differences in expression across conditions (Figure S5B). Interestingly, the promoter accessible region overlapped with key histone marks that may gate chromatin access (Figure 4E). Females also had higher expression of activating chromatin signature H3K27ac (Figure S5C). H3K27ac ChIP analysis in female liver revealed stronger binding of H3K27ac at the *Pnpla3* locus compared to males (Figure S5C and S5D). The balance or activating and repressive histone marks can be orchestrated by chromatin modifying factors Ezh1 and Ezh2. Ezh1 and Ezh2 have been linked with sex-biased regulation and gonadal hormone balance (Lau-Corona et al., 2020; Ma et al., 2017; Roy et al., 2022), which led to us hypothesize that Ezh proteins could contribute to differences in *Pnpla3* expression between the sexes. Double knockout of Ezh1/2 (DKO) in liver resulted reduced differences in *Pnpla3* expression between the sexes (Figure 4M). Collectively, these data suggests that that histone modifying factors that gate promoter regions contribute to sex-differences in regulation of *Pnpla3*.

## Discussion

Our data provides a comprehensive catalog of changes in chromatin dynamics between the sexes under different dietary conditions. Selective gene regulation is strongly influenced by the epigenetic landscape (Kuznetsova et al., 2020; Wilkinson et al., 2023; Zhang et al., 2019). Therefore, mapping nucleosomes dynamics across conditions can be used to infer critical transcriptional regulators and determine their cooperative and hierarchical interactions. Our analysis pointed to a number of unexpected findings. First, chromatin accessibility differed by diet composition was dominantly informed by sex. Second, feeding a western or high fat resulted in greater differences in the chromatin landscape between the sexes. The differences were particularly more pronounced at promotor regions.

Multiple motifs showed common enrichment between male and female liver, but our analysis also identified critical differences in nucleosome binding patterns. For example, females had stronger enrichment of promotor binding factors, like the trimeric complex NFY and ETS transcription factors. These results were corroborated by gene expression integration and align with our observation that female livers had more accessible promoter regions. Male liver showed more extensive nucleosome rearrangements related to nuclear receptor pathways. As expected, we observed differences in established hormonal pathways known to differ between the sexes such as estrogen receptor and androgen receptor.

Integrating chromatin dynamics and gene expression revealed substantial overlap in sex-biased signatures and provides evidence that nucleosome rearrangements inform gene expression changes. However, this was not the case for every transcription factor. For example, LXR-dependent chromatin changes between the sexes were pronounced, yet we observed minimal differences in LXR-dependent gene expression. It is possible there are differences in LXR binding patterns or activity between the sexes but mechanisms beyond RNA biogenesis could dictate steady state transcript levels. For example, change in mRNA stability, export or translation efficiency between the sexes is likely important (Salisbury et al., 2021).

Our unsupervised analysis of sex biased genes highlighted a number of critical factors involved in lipid metabolism and lipid droplet regulation. *PNPLA3* is one of the few genes consistently implicated in MAFLD GWAS (Anstee et al., 2020; Ma et al., 2017; Romeo et al., 2008; Speliotes et al., 2010). Using epidemiologic and molecular studies, recent evidence suggested that regulation of *PNPLA3* by estrogen contributes to the rise in MAFLD in older females (Cherubini et al., 2023). These studies do not consolidate well with known changes in estrogen levels that occur during menopause. Examination of chromatin dynamics at *Pnpla3*, corroborate our overarching findings that changes at promoters appear to underlie differences in hepatic expression of *PNPLA3* between males and females. Integrating our findings with public datasets, we use knockout and overexpression models to show that sex differences in PNPLA3 transcript levels are regulated by male gonadal factors and histone modifiers acting at promoters, rather than estrogen. Our work does not entirely rule out a role for estrogen signaling in *PNPLA3* regulation, but leveraging a ‘chromatin-centric’ platform suggests that other factors may be contributing. Future experiments can more thoroughly interrogate the molecular pathways that regulate sex-differences in *PNPLA3* expression. In summary, we provide a resource that can used to uncover mechanisms of gene regulation in hepatic response to lipid excess between the sexes.

## Limitations of this study

First, while we observed that sex was the pivotal driver of chromatin responses, we do point out that our dietary perturbations were only for 2 weeks. It possible that longer dietary feeding would have shown different results. However, we doubt that this would be the case since important evidence found that genetic control of metabolic response to dietary challenge is most pronounced within the first 2 weeks (Parks et al., 2013), hence the focus on early timepoints. Second, we use the GWAS gene *Pnpla3* as an example of how our resources can inform our understanding of sex or diet-biased regulatory circuits, but the precise molecular details of how gonadal factors regulate *Pnpla3* are yet to be elucidated and will be subject of future work.

## Materials and methods

### Animals and diet

All animal experiments were approved by the UCLA Animal Research Committee and strictly adhered to the guidelines outlined in the Guide for the Care and Use of Laboratory Animals by the National Institutes of Health. The mice used in this study were of the C57BL/6 background and housed in a temperature-controlled room at 22°C with 50–65% humidity, under a 12-hour light/dark cycle in pathogen-free conditions. Wild-type mice (strain 000664) were obtained from The Jackson Laboratory and were fed ad libitum on either a standard chow diet (PicoLab Rodent Diet 20, 5053), Western diet (WD: Research Diets, D12079B), or high-fat diet (HFD: Research Diets, D12492). Four Core Genotypes mice on a C57BL/6 congenic background from a UCLA maintained colony were used as we described previously (Link et al., 2020; Reue and Wiese, 2022). These mice have a deletion of the testis determining gene Sry on the Y chromosome and insertion of Sry transgene onto an autosomal chromosome, which allows for independent segregation of gonad development and sex chromosome complement (Burgoyne and Arnold, 2016). For the ATAC-seq experiments, mice were subjected to diet treatments for 2 weeks, while RNA-seq treatments lasted either 2 or 4 weeks. For the FCG mice experiment, gonadectomy was performed at 10-11 weeks of age under isoflurane anesthesia by aseptic procedures as previously described (Chen et al., 2012). FCG mice were fed ad libitum a high-fat diet for 10 weeks (intact gonad cohort) or 16 weeks (gonadectomized cohort). Prior to sacrifice, mice were fasted for 6 hours, unless specified otherwise, such as in the LXR agonist treatment experiment. Liver tissues were harvested, flash-frozen in liquid nitrogen, and stored at -80°C for subsequent analysis. For ATAC-seq, 3 mice were used for each group. The following number of mice were used for each RNA-seq group: N=3 for female Chow diet 2 weeks (CF2), and female Chow diet 4 weeks (CF4), male chow diet 2 weeks (CM2) and male WD 2 weeks (WM2). N= 4 for Male WD 4 weeks (WM2), female HFD 2 weeks (HF2) and female HFD 4 weeks (HF4) and male HFD 2 weeks (HM2). N= 5 for Male chow diet 4 weeks (CM4), female western diet 2 (WF2) and female western diet 4 weeks (WF4) and male HFD 4 weeks (HM4). For the FCG sequencing, the number of mice used in each group is specified in the figure legends.

### Adeno-associated virus (AAV) transduction

For AAV-mediated Esr1 overexpression, eight-week-old male or female C57BL/6J wild-type mice received intravenous injections of AAV8.TBG.GFP or AAV8.TBG.ESRa (purchased from VectorBiolabs) at a dose of 1 × 10^11^ GC per mouse, administered via the lateral tail vein. One week after AAV transduction, mice were placed on a HFD for 5 weeks. Mice were sacrificed following a 6-hour fasting period. Five mice were used for each group.

### LXR agonist treatment

The LXR agonist treatment experiments in mice were conducted following the protocol described in previous studies (Bideyan et al., 2022; Sallam et al., 2016). In brief, 9-week-old mice were administered GW3965 (Sigma) at a dose of 40 mg/ kg, given via oral gavage 17 hours and again 8 hours before sacrifice. Mice were fasted for 4 hours prior to sacrifice. GW3965 was dissolved in dimethyl sulfoxide (DMSO, Sigma) and delivered in canola oil (Sigma). DMSO was used as the vehicle control. Eight mice were used for “Female Vehicle” group, “Male Vehicle” group and “Male GW3965” group, seven mice were used for “Female GW3965” group.

### RNA extraction and qRT-PCR

Total RNA was extracted using TRIzol reagent (Invitrogen). RNA was reverse transcribed using a homemade reverse transcriptase as described in our previous publication (Sallam et al., 2018). Real-time PCR (qRT-PCR) was used to quantify cDNA, utilizing SYBR Green Master Mix (Diagenode) on a BioRad Real-Time PCR instrument. The primers for amplifying *Pnpla3* were sourced from a previous publication (Smagris et al., 2015). Primers amplifying *Polg1* was from our previous publication (Zhou et al., 2020). Gene expression level was normalized to the housekeeping gene 36B4 or as indicated in Figure Legends.

### Immunoblot analysis

Proteins were extracted using RIPA lysis buffer (Boston Bioproducts) supplemented with a complete protease inhibitor cocktail (Roche). The protein samples were resolved on a 4–12% NuPAGE Bis-Tris Gel (Invitrogen), transferred to a Hybond ECL membrane (GE Healthcare), and then probed with primary antibodies. The following antibodies were used: anti-PDI (Cell Signaling Technology, 3501, 1:1000 dilution) and anti-Estrogen Receptor α (Sigma-Aldrich, 06-935, 1:1000 dilution).

### ATAC-Sequencing and ATAC-Sequencing analysis

ATAC-sequencing (ATAC-Seq) using frozen liver samples were performed using our previous optimized protocol (Zhang et al., 2020). The number of nuclei in each sample was counted using Trypan Blue (Thermo Fisher) and a hemocytometer, and 75,000 nuclei were extracted and centrifuged at 500 g for 5 min at 4°C and used for the transposition reaction. ATAC-Seq libraries were constructed using the Nextera Tn5 Transposase and DNA library preparation kit (Illumina) as described (Buenrostro et al., 2015). We performed size selection usign AMPure XP magnetic beads. Libraries were sequenced by NovaSeq 6000 (Read Length: 2× 100) at the Broad Stem Cell Research Center Sequencing Core (BSCRC) at UCLA.

Sequenced reads from each sample were individually aligned to the mouse genome (mm9, NCBIv37) using Bowtie2 (Langmead and Salzberg, 2012). Duplicated reads or those that mapped to the mitochondrial genome or aligned to unmapped contiguous regions were removed. MACS2 (Zhang et al., 2008) was applied for peak calling for each sample using the option: –nomodel –keep-dup all -q 0.01 –llocal 10000. We merged the reproducible peak sites in each sample and obtained RPKM for each sample by SeqMonk (Babraham Bioinformatics). Bedgraph files were generated by HOMER (Heinz et al., 2010) and transformed to TDF by IGV tools and visualized (Robinson et al., 2011). Gene annotation was performed by Homer annotation function using the genome location of peaks. Motif enrichment analysis was performed by HOMER Motif Analysis (Heinz et al., 2010; Thorvaldsdottir et al., 2013). Promoter specific motif analysis was performed for Figure 2 and global motif analysis was performed for Figure 3. PCA was performed with R package PCAtools (Blighe K, 2024). PCA plots were generated using the mean values from each group (Figure 1C) or the individual values from the samples in each group (Figure S1C), respectively. Peaks including annotation were provided in supplemental Table S1.

### RNA-Sequencing and RNA-Sequencing analysis

RNA sequencing (RNA-Seq) was performed as we previously described (Salisbury et al., 2021). Briefly, total RNA for RNA-sequencing was extracted using TRIzol reagent (Invitrogen) and purified by Qiagen RNeasy Mini Kit (Qiagen). RNA concentration was measured by Nanodrop and sequenced at UCLA Technology Center for Genomics and Bioinformatics (TCGB) using Hiseq3000 (Read Length: 1× 50 bp). RNA-seq analysis was performed as described in our previous publication (Salisbury et al., 2021). Each sample was processed independently, and RPKM values were calculated for each sample individually. Differential expressed genes (DEGs) in diets challenging experiments were defined as: fold change [(WD or HFD)/Chow] > 1.5 & P-value < 0.05 (for up-regulated genes), and fold change [Chow/ (WD or HFD)] > 1.5 & P-value < 0.05 (for down-regulated genes). Statistics of DEGs were provided in supplemental Table S2. Ontologies and Transcription analysis were performed by EnrichR (Chen et al., 2013; Kuleshov et al., 2016; Xie et al., 2021), or Metascape (Zhou et al., 2019) as described in the results or Figure legends. Venn diagrams were generated by Venny (Oliveros, 2007-2015) or Venn diagram online tool from Bioinformatics & Evolutionary Genomics (https://bioinformatics.psb.ugent.be/webtools/Venn/). Heatmaps were generated by Morpheus (https://software.broadinstitute.org/morpheus) or ClustVis (Metsalu and Vilo, 2015). For the liver RNA-sequencing data obtained from public datasets, we utilized the processed data generated by Lau-Corna et al. (Lau-Corona et al., 2020). Related to Figure 4, sex-specific gene expression was compared by calculating the ratio of RPKM values: “Control female” RPKM divided by “Control male” RPKM. To identify female- or male-biased DEGs, we applied the following criteria: 1). Female-biased DEGs: Fold change (FC) of WD/Chow > 1.2 at both 2 and 4 weeks in females, while FC (WD/Chow) ⩽ 1 at least at one time point in males; or FC (Chow/WD) > 1.2 at both 2 and 4 weeks in females, with FC (Chow/WD) ⩽ 1 at least at one time point in males. 2). Male-biased DEGs: FC (WD/Chow) > 1.2 at both 2 and 4 weeks in males, while FC (WD/Chow) ⩽ 1 at least at one time point in females; or FC (Chow/WD) > 1.2 at both 2 and 4 weeks in males, with FC (Chow/WD) ⩽ 1 at least at one time point in females. For the RNA-Sequencing analysis of FCG model, reads were aligned to GRCm39 reference mouse genome and read count per gene quantified with STAR (version 2.7.10a). DEG analysis was performed with DESeq2 (version 1.36) with contrast tests with reduced models to evaluate effect of each variable: gonadal effect (mice with intact ovaries vs. mice with intact testes), chromosomal effect (mice with XX chromosomes vs. mice with XY chromosomes), ovariectomy effect (ovariectomized mice vs. mice with intact ovaries), and castration effect (castrated mice vs. mice with intact testes).

### ChIP-Sequencing analysis

ChIP-Sequencing (ChIP-seq) from public datasets are as listed in figure legends. We reanalyzed the ChIP-seq using downloaded fastQ files. Reads were aligned to the mouse genome mm9 (NCBIv37/mm9) using Bowtie2 (Langmead and Salzberg, 2012). MACS2 (Zhang et al., 2008) was applied for peak calling. To visualize the peaks, HOMER software was applied to generate bedgraph files, which were then transformed to TDF and visualized by IGV tools (Robinson et al., 2011; Thorvaldsdottir et al., 2013). Gene annotation was performed by SeqMonk annotated to “mRNA” (Babraham Bioinformatics).

## Data availability

The ATAC-sequencing and RNA-sequencing datasets have been deposited to the National Center for Biotechnology Information and are available at accession number: ATAC-seq: (Pending). RNA-sequencing for diets treatment: (Pending). RNA-Sequencing for FCG mouse model: (Pending).

## Acknowledgments

This work was supported by National Institute of Health (NIH) grants (DK118086, HL139549, HL149766). We would like to thank the UCLA Division of Laboratory Animal Medicine (DLAM) for assistance with animal studies. We thank the staffs at the TCGB and BSCRC of University of California Los Angeles (UCLA) for their services and assistance. We thank members of the Sallam Lab for helpful comments and suggestions.

## Footnotes

This article is accompanied by supplemental material.

---

**Figure S1.** ATAC-seq in mice liver (Related to Figure 1). (A) Total read count of each ATAC-seq sample. Values shown as Log_10_(total read count). (B) Total alignment rate from each sample was calculated by bowtie2 using mm9 as reference genome. (C) PCA plots of chromatin accessibility of liver samples generated using individual peak score from each sample (n=122,587). FChow: female mice with normal chow diet feeding, FWD: female mice with western diet feeding, FHFD: female mice with high fat diet feeding, MChow: male mice with normal chow diet feeding, MWD: male mice with western diet feeding, MHFD: male mice with high fat diet feeding. (D) and (E) Normalized ATAC-seq signal intensity for differential peaks of HFD compared to chow ± 3000 nt (D) or ± 500 nt (E) from TSS. (F). Number of differential female and male peaks in Promoter-TSS region according to Homer annotation.

**Figure S2.** Promoter-binding factors are enriched in female liver (Related to Figure 2). (A) Heatmap of motif accessibility across promoter specific peaks ranked based on accessibility difference between HFD and chow diet samples or WD and chow diet samples, separated to 6 bins for each group. Female and male are calculated separately. Motif analysis performed by Homer. The numbers within the colored heatmap represent the –Log_10_ (P-value) for each motif. Only motifs with a maximum –Log_10_ (P-value) greater than 5 are included. (B) Heatmap showing maximum –Log_10_ (P-value) of male or female enriched motifs from Figure S2A. Cluster A represents female-preferred transcription factors. Cluster B represents male-preferred transcription factors. Values indicated –Log_10_ (P-value). (C) Venn diagram showing the overlap of the top 10 enriched TFs from each group. (D) Bar plot showing the maximum –Log_10_ (P-value) for females (F ^max^) minus that for males (M ^max^).

**Figure S3.** Sex-specific hepatic gene expression profiles exhibit different motif preferences (Related to Figure 3). (A) Heatmap showing data of all the genes detected by RNA-seq in mice liver samples. (B) Differentially expressed genes (DEGs) in male or female with 2 weeks or 4 weeks WD feeding. (C) DEGs in male or female with 2 weeks or 4 weeks HFD feeding. (D) Venn diagram showing female and male common DEGs and specific DEGs. (E) Metascape analysis of male and female common DEGs. Top 5 terms shown. (F-I). Top 10 Metascape terms shown for female or male-specific up or down regulated genes. (J) Motifs enrichment prediction of male-biased or female-biased genes in Figure 3F. (K) Genes from Figure 3F matching to DEGs identified by RNA-seq using FCG mouse model. P-value< 0.05 was considered as significant effect. (L) Venn diagram exhibiting overlap of diets induced sex difference genes from Figure 3F and DEGs identified by RNA-seq using FCG mouse model.

**Figure S4.** Gene expression from human liver corroborates analysis from murine studies (Related to Figure 4). (A) Heatmap was generated based on TPM of genes from Figure 4A in human livers. TPMs were obtained from GETx Portal. Genes from Figure 4A with no related human genes, or with expression level of “0” were taken out. (B and C) Venn diagram showing the overlap of female (B) or male (C) diets induced DEGs from Figure 4A and DETx genes. (D) Expression of human PNPLA3 in top 10 tissues. Data from GETx Portal. (E) Venn diagram showing the overlap of DEGs from Figure 4F and liver expressed sex-biased genes from Lau-Corona et al., 2020. (F) Fold-change of sex-biased DEGs from Lau-Corona et al., 2020. Fold changes shown as Males/Females (control mice from Lau-Corona et al., 2020). (G) LXR target gene expression based on RNA-seq from Figure 3A. CF2: chow diet fed female mice for 2 weeks. WF2: WD fed female mice for 2 weeks. CF4: chow diet fed female mice for 4 weeks. WF4: WD fed female mice for 4 weeks. CM2: chow diet fed male mice for 2 weeks. WM2: WD fed male mice for 2 weeks. CM4: chow diet fed male mice for 4 weeks. WM4: WD fed male mice for 4 weeks. (H) Expression of LXR target genes detected by qPCR from liver. N=8 per group. P-value was calculated by unpaired t-test. *: P-value< 0.05; **: P< 0.01; ***: P< 0.001; ns: not significant.

**Figure S5.** Chromatin architecture and regulation of *Pnpla3* (Related to Figure. 4). (A) Gene expression of *Polg1* detected by qPCR from livers of mice from Figure 4F, shown as relative to male or female AAV-TBG-GFP values respectively. N= 5; * P< 0.05 by t-test. (B) PROMO predicted TFs binding sites around TSS of *Pnpla3*, as indicated in the red box in Figure 4E. “Level” revealed=s the density of binding sites of TFs, higher “level” indicated the higher density of TFs binding. (C) H3K27ac ChIP-seq peaks from public datasets under GEO#: SRR6756579 (female Rep 1), SRR6756580 (female Rep 2), SRR6756581 (female Rep 3), SRR6756582 (female Rep 4), SRR6756571 (male Rep 1), SRR6756572 (male Rep2), SRR6756573 (male Rep3) and SRR6756574 (male Rep 4). (D) Fold change calculated based on ChIP-seq enrichment of H3K27ac performed in male or female liver (from Lau-Corona et al., 2020). Gene annotation was performed by Seqmonk annotated to mRNA.

