## Supplementary figures and images for "A genome-wide ATLAS of liver chromatin accessibility reveals that sex dictates diet-induced nucleosome dynamics"

### Supplemental Figures

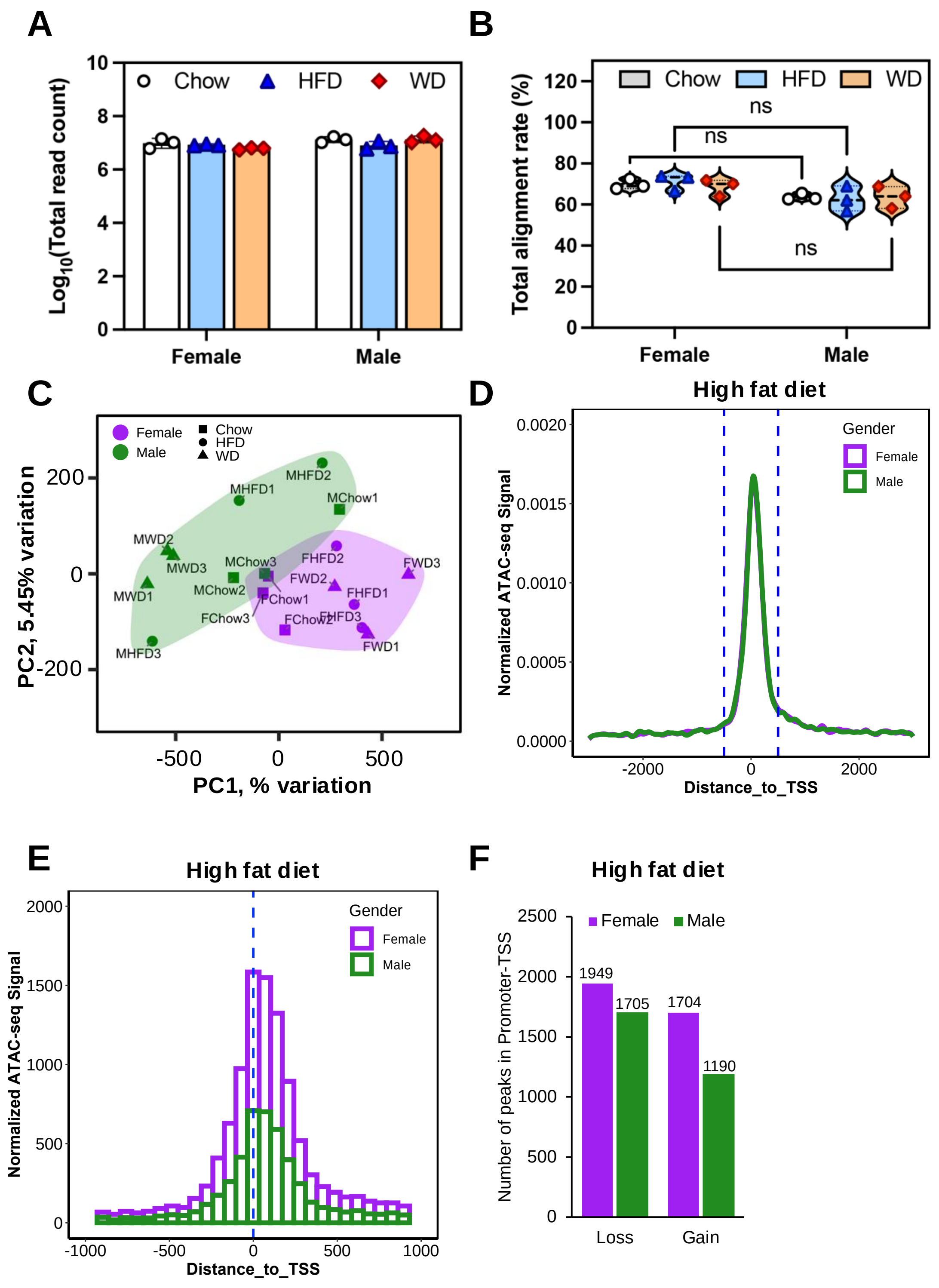

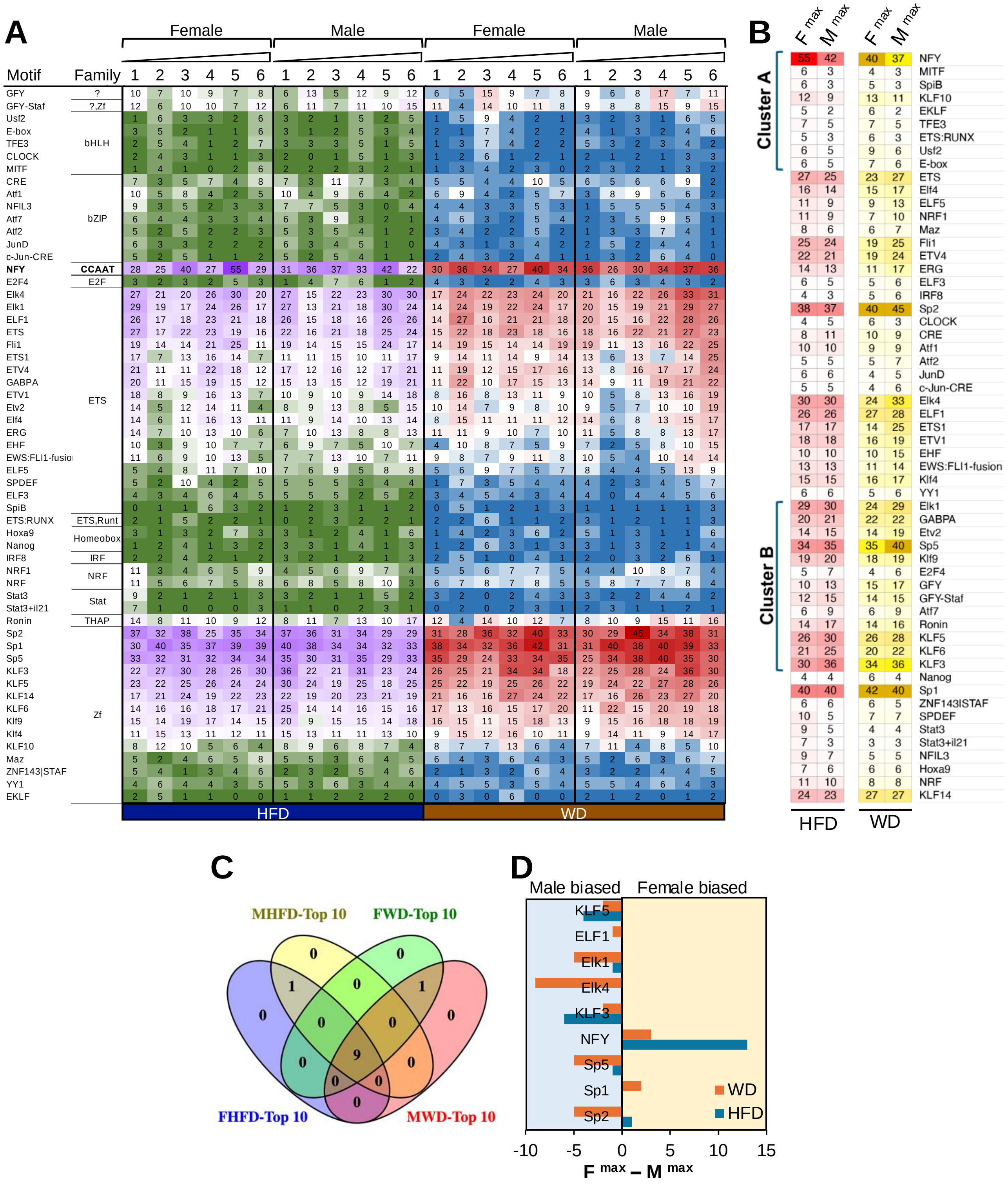

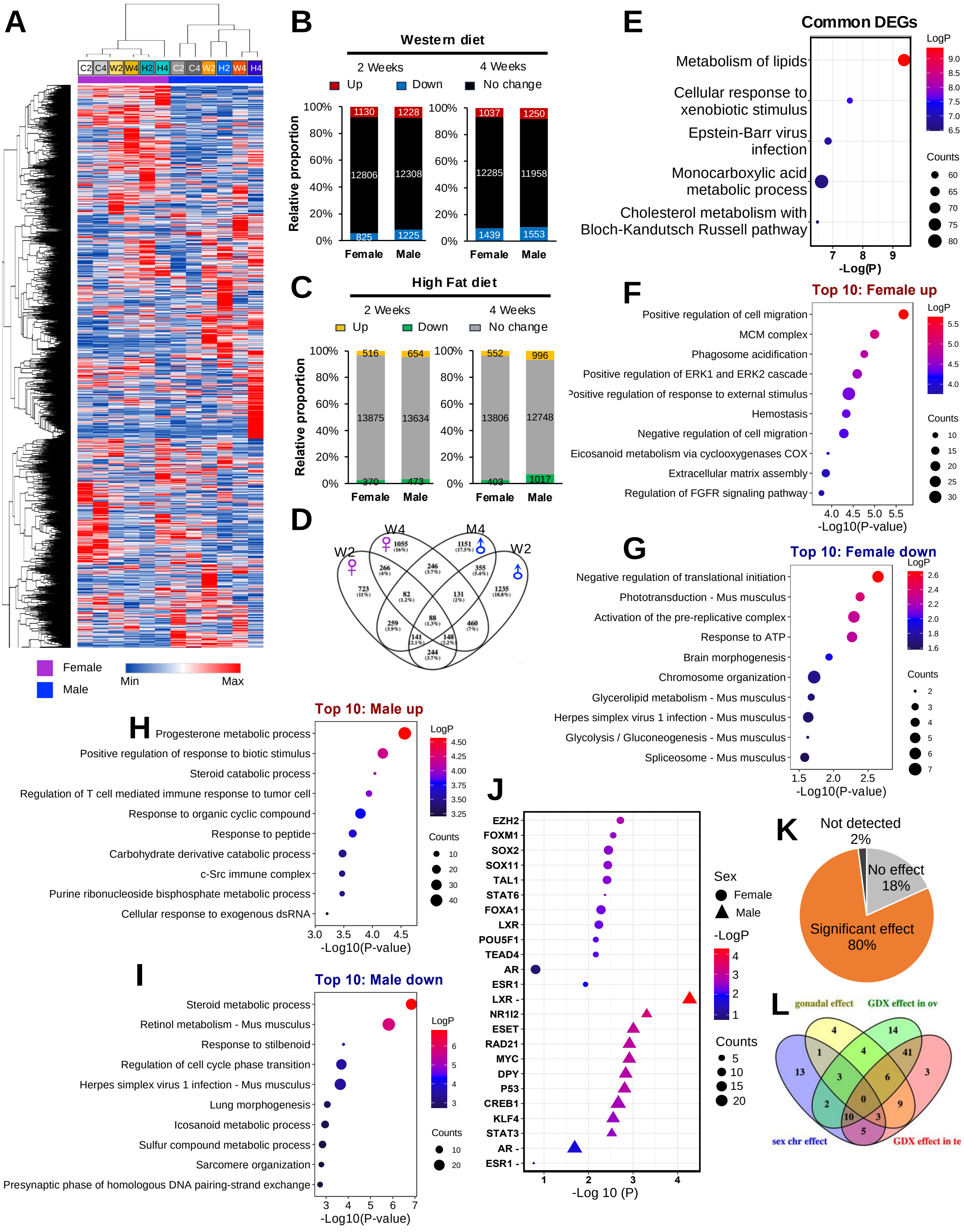

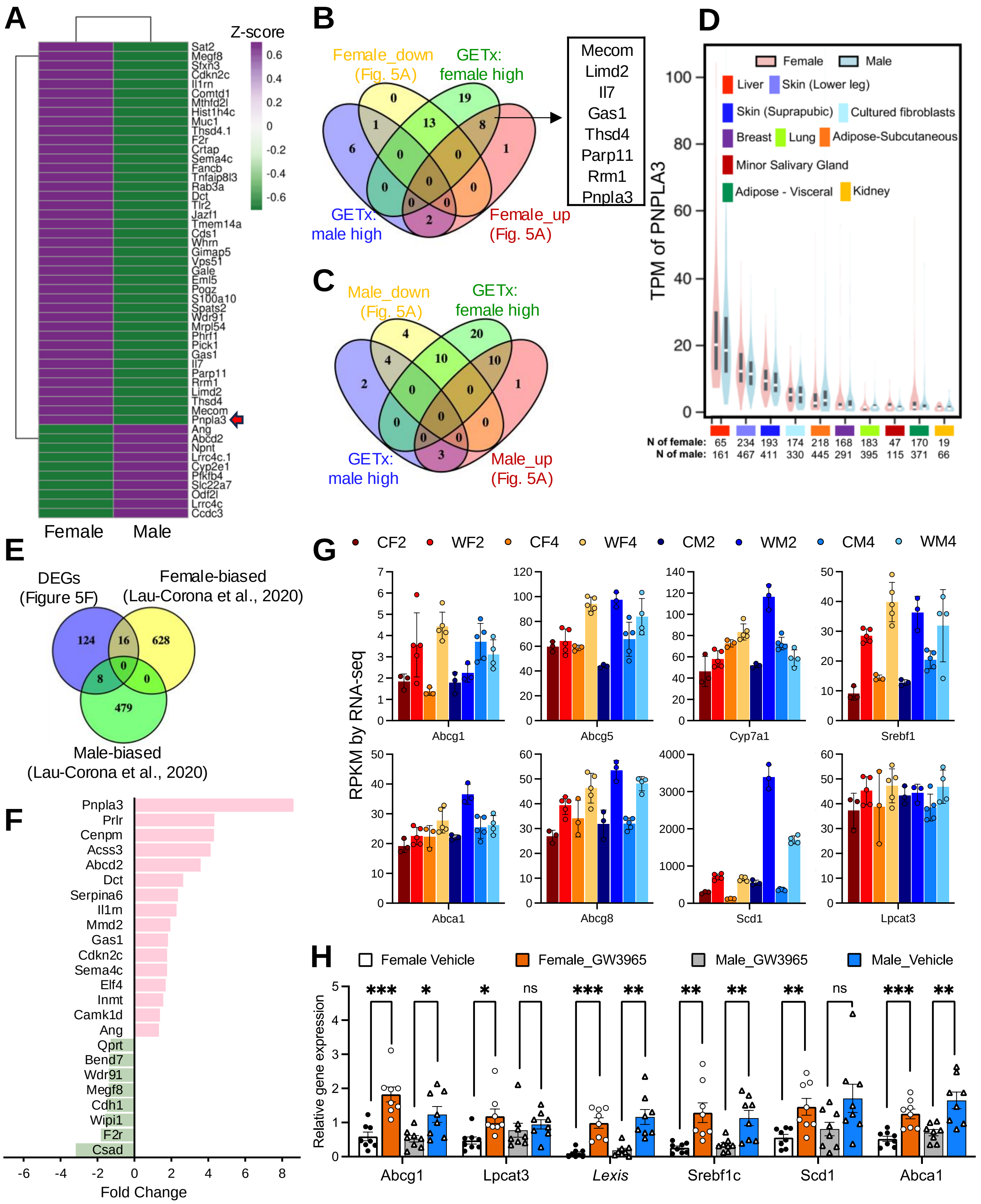

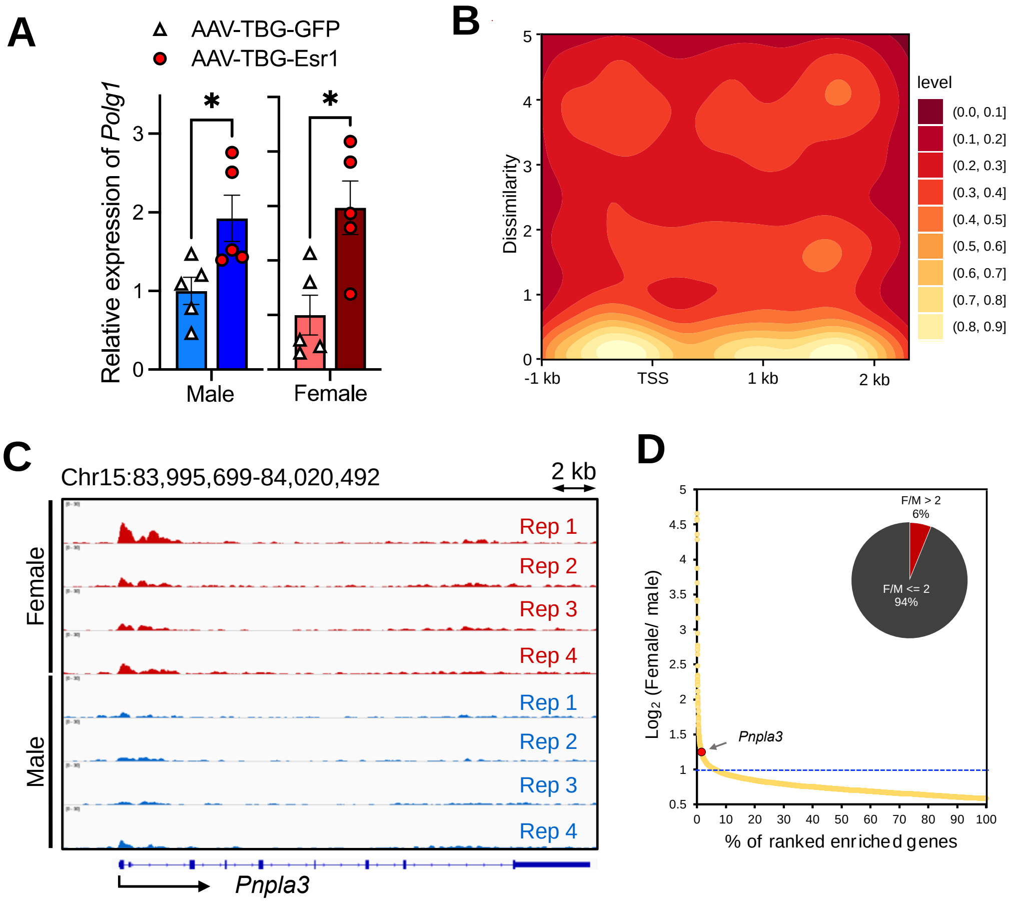
